# A cognitive process model captures near-optimal confidence-guided waiting in rats

**DOI:** 10.1101/2024.06.07.597954

**Authors:** J Tyler Boyd-Meredith, Alex T Piet, Chuck D Kopec, Carlos D Brody

**Author notes:** correspondence should be addressed to: Carlos D Brody.

## Abstract

Rational decision-makers invest more time pursuing rewards they are more confident they will eventually receive. A series of studies have therefore used willingness to wait for delayed rewards as a proxy for decision confidence. However, interpretation of waiting behavior is limited because it is unclear how environmental statistics influence optimal waiting, and how sources of internal variability influence subjects’ behavior. We trained rats to perform a confidence-guided waiting task, and derived expressions for optimal waiting that make relevant environmental statistics explicit, including travel time incurred traveling from one reward opportunity to another. We found that rats waited longer than fully optimal agents, but that their behavior was closely matched by optimal agents with travel times constrained to match their own. We developed a process model describing the decision to stop waiting as an accumulation to bound process, which allowed us to compare the effects of multiple sources of internal variability on waiting. Surprisingly, although mean wait times grew with confidence, variability did not, inconsistent with scalar invariant timing, and best explained by variability in the stopping bound. Our results describe a tractable process model that can capture the influence of environmental statistics and internal sources of variability on subjects’ decision process during confidence-guided waiting.

## Introduction

A decision maker’s estimate of the probability that a decision is correct given the evidence is referred to as decision confidence^1,2^. Confidence is critical for learning improvements in decision policy in response to feedback^3^, for deciding whether to act or gather more information^4^, and for determining how to long to wait for an expected outcome before seeking reward elsewhere^5^. However, study of the neural underpinnings of confidence has been limited by the difficulty of measuring confidence in animal subjects.

Recent work^6,5^ has developed a promising assay of decision confidence by asking how long subjects are willing to wait for a rewarding outcome after a decision before moving on to another reward opportunity. This temporal post-decision wagering paradigm has been used to study confidence in olfactory^5,7^, auditory^7^, visual^8^, and mnemonic^9^ decisions, and has been used to study hallucinations^10^. Temporal post-decision wagers have also been used to probe learning about environmental reward statistics^11^.

The premise of these confidence-guided waiting studies is that reward-rate-maximizing (“optimal”) agents are willing to wait longer for delayed rewards when they are more confident in their decisions. Modulations in willingness to wait are therefore understood to reflect variations in decision confidence. However, optimal waiting behavior also depends on the opportunity cost of opting to continue waiting for reward rather than starting a new trial, which is set by the maximum achievable environmental reward rate^5,12,13,14,15^. Previous work on this task did not explicitly define the environmental reward rate and therefore cannot specify how to choose the optimal average willingness to wait in a given environment, which is the first-order statistic that needs to be optimized in order to maximize reward rate. A complete account of optimal behavior in this task requires a definition of the reward rate that specifies all relevant environmental statistics affecting the environmental reward rate. Without such an account, it is not possible to determine whether animal subjects performing this task are behaving optimally.

Here, we trained rats to perform an auditory evidence accumulation task^16^ requiring binary decisions followed by a temporal wager^5^. We developed an expression for the reward rate in the task, which allowed us to find the reward-rate-maximizing waiting policy. Doing so made explicit a key environmental statistic: the travel time that is incurred when moving on from one reward opportunity to the next. We found that rats performing the task spent longer waiting for rewards than optimal agents who maximized reward rate on matched datasets. This finding was consistent with the observation of “overharvesting” in foraging studies^17,14^. However, when we measured each individual subject’s travel times and treated these as constraints on agents’ behavior, the rats’ overall average willingness to wait was not different from the optimal agents’, suggesting that their waiting behavior was approximately optimal.

In addition to finding near-optimal overall average waiting behavior, the rats’ also showed modulation of wait times by decision confidence, as has been seen previously^5,7,8,10^, consistent with optimal agents. However, it is not clear how the near-optimal behavior we observed can be executed algorithmically in the brain. Nor is it clear how that decision might evolve in time and be influenced by sources of internal variability other than confidence. To develop a candidate model of the waiting decision process, we used the sequential probability ratio test (SPRT) to derive a decision variable that could achieve optimal waiting via an accumulation to bound process, as is often used in decision-making tasks^18,19,20^. The decision variable was initialized at a point encoding the decision confidence and evolved with a linear drift toward a single fixed bound that encoded the estimate of the environmental reward rate. The drift in this model came from the observation that as time elapses without reward after a decision, the odds that the trial will be rewarded eventually decrease. Under the model, waiting continues until the moment that the decision variable crosses the bound at which point the current trial is abandoned.

The process model allowed us to compare various mechanistic sources of noise that might effect the decision process^16^. We considered variability in the drift rate that would produce the property of scale invariance often observed in the literature on timing judgments^21,22,23^, whereby the standard deviation of timing judgments grows proportionally to the interval being timed. We also considered diffusion noise that would corrupt the decision variable in each time step and cause the standard deviation of timing judgments to scale with the square root of the interval being timed, as in the drift diffusion model^24^. Finally, we considered variability in the setting of the bound, which would cause variability in timing judgments to be constant across all intervals being timed. Surprisingly, we found that, whereas scale invariant timing noise had been assumed to dominate during this task^5^, the data was most consistent with variability in the bound. We speculate that the dominance of this source of variability may arise from continual learning of the bound setting based on a recency weighted average of the reward history, as has been observed in previous studies^14,11^ and is used in models of foraging as evidence accumulation^25^.

Our results lay out a more complete theory of optimal behavior in temporal post-decision wagering tasks and present a process model for estimating the moment-to-moment cognitive state of subjects during the waiting decision process. Taken together, these contributions increase the interpretive value of post-decision temporal wagers for studies of confidence and learning.

## Results

### Evidence accumulation task with confidence-guided waiting

We trained rats (n=16) to perform an auditory evidence accumulation task^16^ with randomly delayed reward delivery (Fig. 1a), as in Lak et al.^5^. The task requires two decisions of interest. First, the animal should decide which of two reward ports is more likely to provide a water reward given the auditory stimulus. Then, the animal must decide how long to wait for reward to be delivered before moving on to the next trial. In principle, this decision could be made at the time of the port choice, but may also be characterized as a series of decisions to wait at the chosen port for another timestep or abandon the port and move on.

**Figure 1:**
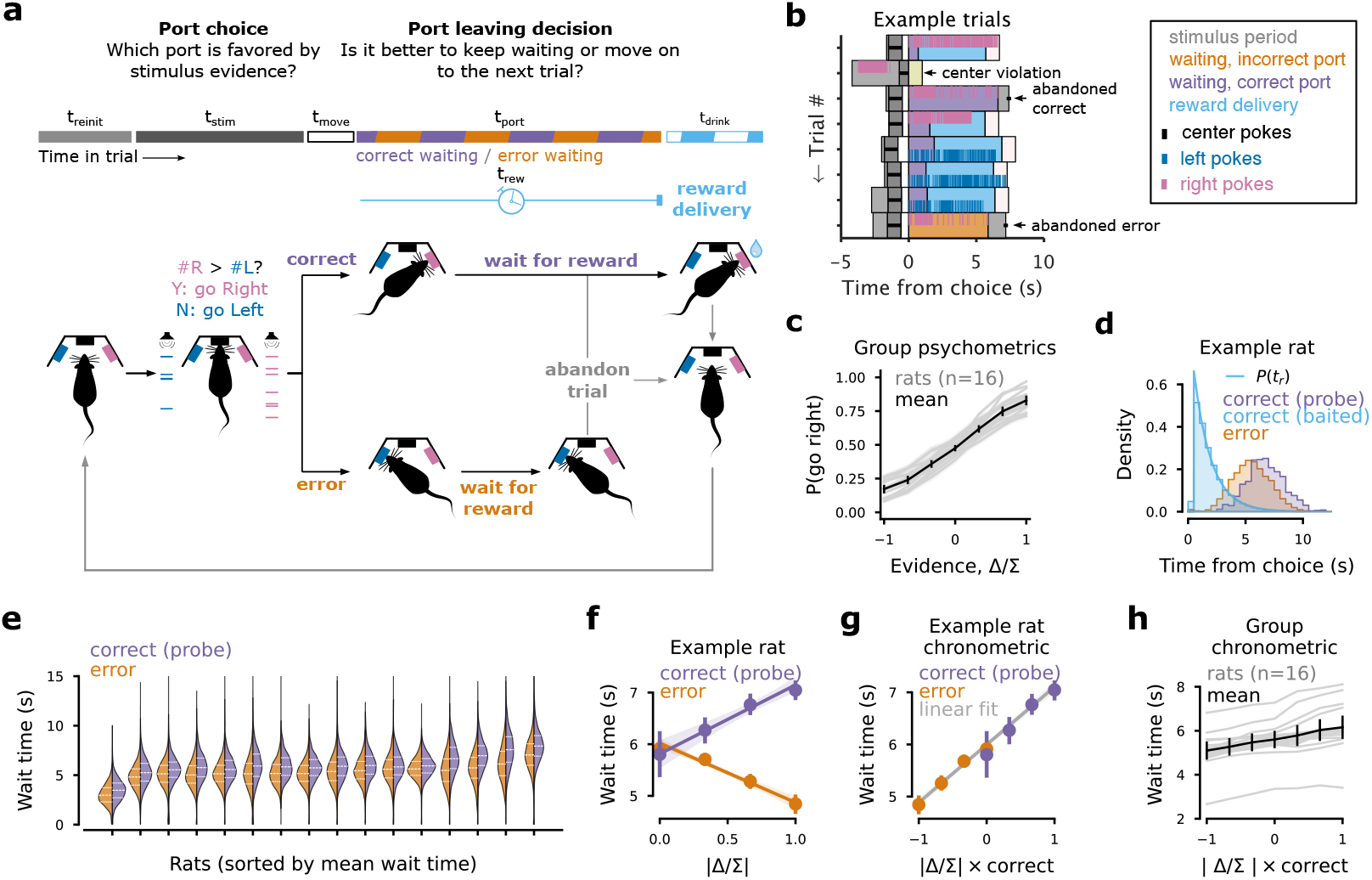
Evidence accumulation task with delayed rewards. (A) Schematic of the task structure. Rats first make a port choice based on an auditory stimulus. On non-probe trials, if the port choice is correct, a reward is scheduled for delivery. The rat waits at the chosen port until either the reward is delivered or the rat decides to abandon the current trial and move on to the next. (B) Example trials from an example rat aligned to the time of the port choice. Correct port choices (purple bars) often lead to rewards (cyan bars), but sometimes the animal abandons the trial and center pokes to start a new trial before reward is delivered (an example is annotated). Error trials (orange bar) are never signalled and the rat eventually has to abandon the trial (an example is annotated). If the animal fails to hold it’s nose in the center port during the stimulus period, the trial is considered a “center violation” (an example is annotated), increasing the time to the next possibly rewarded trial. (C) Probability of choosing the rightward reward port is plotted as a function of the evidence, operationalized as the click difference (#*R* − #*L*), normalized by the total number of clicks (#*R* + #*L*), denoted Δ*/*Σ. Each rat is shown as a gray trace and the average (with 95% confidence intervals) of the gray curves is shown in black. (D) Wait time distributions for an example rat conditioned on whether the decision was incorrect (shaded orange), correct with a baited reward (shaded cyan), or a correct probe trial (shaded purple). The reward delay distribution used to draw reward delivery time on non-probe trials is underlaid (cyan trace). (E) Violin plots showing each rat’s wait times for error trials (orange) and correct probe trials (purple). Medians are plotted as dashed white lines with 25^th^ and 75^th^ as dotted white lines. (F) Mean wait time for an example rat in correct probe trial (purple) and error trials (orange) as a function of the absolute evidence strength, |Δ*/*Σ|. Data are overlaid on linear fits to the correct and error trials separately. Errorbars are bootstrapped 95% confidence intervals. (G) Wait time chronometric curve showing the mean wait times from F plotted as a function of the strength of evidence favoring the chosen option, |Δ*/*Σ|× correct. Data are overlaid on a linear fit to all the data. (H) Wait time chronometric curve for each rat computed as in g (gray traces) along with the mean (with 95% confidence intervals) of all the gray traces (black trace).

### Port choice

Rats performed the task in a chamber containing an array of three nose ports. Rats initiated trials by poking their nose into a central nose port, which triggered stimulus playback. The stimulus consisted of two trains of auditory clicks, generated from two different Poisson rates, played from speakers on either side of the rat’s head. The rat’s task was to listen to the click trains and then, after a “go” cue, report which click train had the larger number of clicks by poking it’s nose into the reward port on the side associated with the higher click rate. Trials where the rat withdrew from the center port before the “go” cue were labeled “center poke violations,” invalidated, and the rat was moved on to the next trial after a brief white noise stimulus.

There were two versions of this task, referred to as the location task and the frequency task. Each rat was trained to perform one of the two tasks (location task, n=9; frequency task, n=7). In the location task, one click train was played from a speaker to the left of the center port and the other click train was played from a speaker to the right of the center port. Rats were rewarded for choosing the port on the same side as the speaker that emitted the greater number of clicks. In the frequency task, the clicks were played in stereo, but the clicks in the two click trains were played at different frequencies. The high frequency click train instructed rightward choices, whereas the low frequency click train instructed leftward choices. The click trains depicted in Fig 1a correspond to the location task. Trial difficulty was controlled by varying the rates of the two click trains.

### Wait time decision

After reporting a decision at the reward port, rats did not receive immediate feedback. Instead, after correct choices, reward delivery was delayed until an experimenter-determined reward time, *t*_*r*_. On a subset of correct trials (mean: 7.1% ±1.6%), rewards were omitted to provide an uncensored report of the rat’s maximum willingness to wait on that trial. We refer to these trials as “probe” trials (but, note that “catch” trials is another common terminology). On error trials, feedback was omitted entirely. Rats were allowed to give up waiting for reward and move on to a new trial at any time after making the port choice by withdrawing from the reward port and poking into the center port. A series of example trials is shown in Figure 1b.

In these experiments, reward delays were drawn from an exponential distribution, so that the density of reward delays was given by

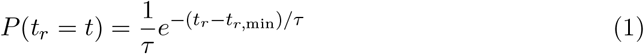

with time constant *τ* = 1.5, and a minimum reward delay *t*_*r*,min_ ∈ (.05, .5). The exponential distribution has a flat hazard rate on the interval *t* ∈ (*t*_*r*,min_, ∞). This means that, given that reward is set to be delivered on a trial, but hasn’t been delivered so far by time *t*_*w*_, the probability of receiving reward in the next time step is constant. We will write the hazard rate of the reward distribution as

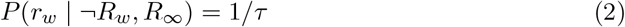

where *r*_*w*_ ≡ *t*_*r*_ ∈ (*t*_*w*_, *t*_*w*_ + *δt*) is used to indicate the event that reward is delivered in the infinitesimal timestep *δt* beginning with time *t*_*w*_, the sum 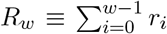 is used to indicate whether reward is baited at some time before *t*_*w*_, and the negation, ¬*R*_*w*_, indicates that no reward is baited before *t*_*w*_. In this notation, *R*_∞_ is used to indicate whether reward is set to be delivered eventually in the trial. The resulting mean reward delay is ⟨*t*_*r*_⟩ ≈ *t*_*r*,min_ + *τ* for trials where reward was baited.

### Rat behavior

All rats included in the study learned to perform the task with at least 60% accuracy (group mean: 74.4% correct trials). We computed each rat’s psychometric function for the port choice (Figure 1c). The psychometric function was defined as the probability of making a rightward choice given the stimulus evidence favoring rightward choice. Stimulus evidence favoring rightward choice is defined as the click difference normalized by the total number of clicks, 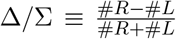, where #*R* represents the number of clicks favoring rightward choice and #*L* represents the number of clicks favoring a leftward choice on a given trial.

We measured time spent waiting for reward at the side port in three trial types of interest: error trials, correct probe trials (where no reward is baited), and correct trials where reward is baited. We excluded trials where the animal took more than 2 seconds to initiate a new trial by center poking after leaving the chosen port. This is a standard criterion used to focus analysis on trials where the animal is engaged in the task^5^. The example rat was willing to wait long enough to receive reward on most correct trials, so the distribution of waiting times on correct trials where reward was baited closely resembles the reward delay distribution (Figure 1d).

On trials where the rat waited long enough to receive reward, the full duration that the rat would have been willing to wait is unknown, because reward delivery censors our observation of the full willingness to wait. We used the probe trials to measure how long rats were willing to wait on correct trials. On both error trials and correct probe trials, rats were willing to wait much longer than the typical reward delays (Fig 1d,e). This held across all rats who learned the task (Fig 1e). Additionally, all rats waited longer at the choice port after correct choices on probe trials than after errors (Fig. 1d,e; *p <* .01, rank-sum test, 16/16 rats), indicating that waiting was guided by an internal estimate of decision accuracy.

To measure the modulation of wait time by the stimulus evidence, we plotted the example rat’s average wait time as a function of the absolute stimulus strength, |Δ*/*Σ|, separately for correct probe trials and error trials (Fig 1f). We expected correct trials with strong signal to be the trials in which the animal has the highest confidence on average. Indeed, wait times were longest for correct trials where the evidence most strongly favored the choice made by the animal, as has been seen previously^5,7,10^. Correspondingly, wait times were shortest in the trials where the evidence most strongly favored the alternative not chosen by the animal. We expect these to be the trials where the animal has the lowest confidence on average.

To create an axis along which both confidence and wait time should increase monotonically, we used the strength of the evidence supporting the option chosen by the rat, |Δ*/*Σ| × correct. This quantity takes positive values when the animal makes a correct choice and negative values when the choice is incorrect. When we plot the example rat’s wait times against this axis, we see a graded increase in wait time as a function of evidence supporting the choice (Fig 1g). The data is overlaid on a linear fit to the data, which has a significantly positive slope (Pearson’s *r* = .22, *p <* .01). We computed wait time as a function of evidence supporting choice for all rats (Fig 1h) and computed linear fits to each rat. All rats had a significant, positive relationship between waiting time and the strength of evidence for the chosen option (*p <* .01 for 16/16 rats). This indicates that all of our rats modulated waiting times by their decision confidence.

### Overall reward rate maximization depends on travel time

Previous work^5^ has shown that in order to maximize reward rate in decision tasks with delayed reward, subjects should be willing to wait longer when they are more confident in their decisions. However, the trial-by-trial modulation of waiting time by confidence alone is not enough to maximize the long term average reward rate. To maximize the long term average reward rate, subjects must also find an appropriate overall average willingness to wait. This value depends on a variety of other environmental statistics that influence the environmental reward rate. However, previous work has not developed an explicit expression for the environmental reward rate in the task, so it has not been possible to test whether rats learn this first-order optimization of overall wait time. In another study of rats performing confidence-guided waiting for delayed rewards, Stolyarova et al.^8^ noted that their rats’ overall wait times were long relative to the average time of reward delivery, as is true in our rats. The authors interpreted this observation as likely being a deviation from optimality in rat behavior. This would be consistent with previous studies in human^14^ and animal subjects^17,26^ which report a bias, referred to as “overharvesting,” toward spending more time than would be optimal on a given reward opportunity before moving on to the next. Here, we develop a definition of the reward rate that makes all relevant environmental statistics explicit. We can then determine the optimal average willingness to wait for a given environment, making it possible to test whether subjects achieve optimal behavior in the task.

To develop an expression for reward rate in our task, we make use of optimal foraging theory^12,13^, which describes the optimal time an agent should spend in a series of “patches” containing depleting, continuous rewards before traveling to the next patch. In each trial, we think of the rat’s nose poke into the chosen reward port as an entry into a “patch.” We refer to the time spent at the port as *t*_port_. We refer to the elapsed time between leaving the reward port on a given trial and entering a reward port on the next trial as “travel time” and note it in equations as *t*_0_ (Fig. 2a). In this task, the travel time includes the period of trial reinitiation from leaving the reward port on the previous trial to entering the center port on the current trial (labeled *t*_reinit_ in Fig 2a), plus the stimulus period during which the rat hears the stimulus to inform the port choice (labeled *t*_stim_ in Fig 2a), and the time it takes to move from the center port to the chosen reward port (labeled *t*_move_ in Fig 2a). Unlike in the classical foraging theory where reward is continuous and the agent has perfect information about the patch identity, rewards in our task are stochastic, limited to at most one per trial, and the subject has only partial information about whether it is in a rewarded or unrewarded patch. The optimal strategy for such a task has been described for environments where rewarded and unrewarded patches occur with equal probability^15^. We generalize that theory to arbitrary initial probability of being in the rewarded patch.

**Figure 2:**
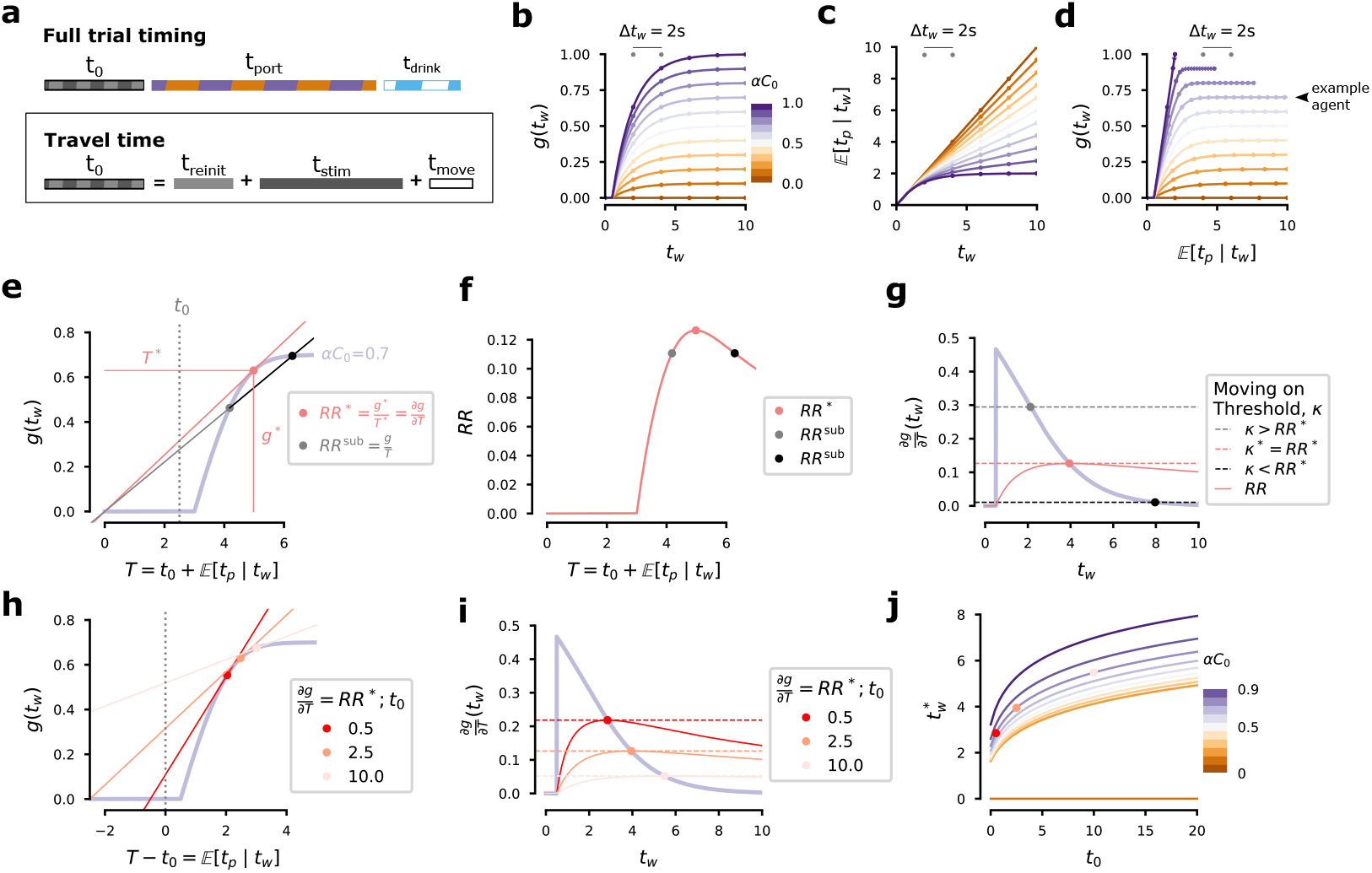
Across trial reward maximization depends on travel time. (A) Task timing can be broken into travel time, *t*_0_, time at the port, *t*_port_, and time spent drinking reward, *t*_drink_ (top). Travel time includes all periods between the end of the waiting/drinking period and the start of the next period of waiting, including trial reinitiation time, *t*_reinit_, stimulus playback time, *t*_stim_, and movement time, *t*_move_ (bottom). (B) Expected reward per trial, *g*(*t*_*w*_), if willing to wait for time *t*_*w*_. Colormap shows probability of eventual reward, *αC*_0_. Points indicate 2s increments of *t*_*w*_. (C) Expected time spent at the port per trial, 𝔼 [*t*_port_| *t*_*w*_], plotted as in B. (D) Expected reward in a trial, *g*(*t*_*w*_), as a function of expected time at the port, 𝔼 [*t*_port_ | *t*_*w*_], plotted as in B and C. The effect of an additional 2s increment in *t*_*w*_ now depends on *t*_*w*_ and *αC*_0_. (E-J) Consider reward maximization for an example agent with *αC*_0_ = .67 on every trial. (E) Expected reward per trial, *g*(*t*_*w*_), plotted as a function of trial time, *T* (*t*_*w*_) = *t*_0_ + 𝔼 [*t*_port_ | *t*_*w*_] (solid purple trace). The reward rate is 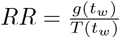 and is maximized when 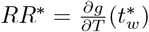 (red point) at 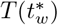. The red trace from the origin through this point has the highest slope of any line from the origin through the purple trace. All other values of *T* (*t*_*w*_) are suboptimal (e.g., gray and black points achieve the reward rate *RR*^sub^, which is the slope of the gray and black traces). Here, *t*_0_ is set to 2.5s. (F) The reward rate for the example agent is plotted as a function of *T* (*t*_*w*_) (dashed red trace) with the maximum reward rate marked (red point) along with the reward rate achieved if not willing to wait long enough (gray point) or willing to wait too long (black point). (G) The instantaneous reward expectation within a trial after time *t*_*w*_ passes without receiving reward, 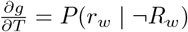 is plotted as a function of *t*_*w*_ (solid purple trace). The session reward rate from F is shown for comparison (solid red trace). Reward rate is maximized if the agent sets a moving on threshold, *κ* = *RR*^*∗*^ (dashed red trace). Suboptimal reward is achieved when *κ* is not set to *RR*^*∗*^ (e.g., gray and back traces). (H) Probability of reward, *g*(*t*_*w*_) = *P* (*R*_*w*_), plotted as a function of expected port time, *T* (*t*_*w*_) − *t*_0_ = 𝔼 [*t*_port_ | *t*_*w*_] (rather than as a function of *T* as in E). Reward maximizing solutions are marked for three values of *t*_0_, including the same from E, one (darkest red trace) smaller, and one larger (lightest red trace). (I) Instantaneous reward expectation plotted as in G with the reward rates and optimal settings of *κ* for the three levels of *t*_0_ used in H. (J) Optimal wait time, 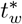, as a function of travel time, *t*_0_, for all levels of *αC*_0_ used in panels b-d, except *αC*_0_ = 1, which corresponds to 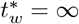. Example levels of *t*_0_ are marked for the example level of *αC*_0_ (red points).

The partial information about patch type comes from the agent’s decision confidence, an estimate of the probability that the port choice was correct at the time of the decision given the available perceptual evidence, which we write as

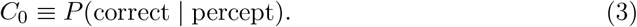

If the agent believes that all correct choices are rewarded, then their initial estimate of the probability that the choice will eventually be rewarded is *C*_0_. If the agent believes that correct choices are only rewarded in some fraction, *α*, of non-probe trials, then the probability that the choice will be rewarded eventually is

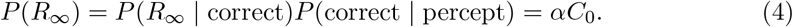

Later, we will see that the agent’s posterior belief about whether the port choice will be rewarded (rewarded or unrewarded) falls over time.

We will simplify the expression for reward rate as a function of the subject’s willingness to wait for reward across trials by beginning with the case of an agent who has no trial to trial variation in decision confidence (i.e., *P* (correct | percept) = *P* (correct)). This agent should therefore be willing to wait the same amount of time, *t*_*w*_, on every trial. We can write the expected overall reward rate for an agent willing to wait until time *t*_*w*_ as the ratio of expected reward per trial, *g*(*t*_*w*_), and expected time per trial, *T*_total_(*t*_*w*_):

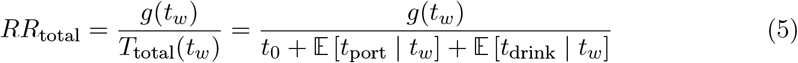

where *t*_0_ is the “travel” time between leaving the chosen side port on one trial and nose poking at a side port on the next trial, 𝔼 [*t*_port_ | *t*_*w*_] is the expected time spent at the side port, and 𝔼 [*t*_drink_|*t*_*w*_] is the expected time spent consuming reward. While 𝔼 [*t*_drink_ | *t*_*w*_] affects the overall reward rate, it can be ignored for the reward maximization process (see Supplemental Information for derivation).

Both expected reward on each trial, *g*(*t*_*w*_), and expected time at the port, 𝔼 [*t*_port_ |*t*_*w*_], depend implicitly on the probability that reward will be delivered eventually if the agent waits long enough (*P* (*R*_∞_); equation 4). Expected reward per trial rises exponentially as a function of willingness to wait toward an asymptote at *αC*_0_ (Figure 2b; see Supplemental Information for mathematical details). Expected time at the port per trial increases as a function of willingness to wait, but does not asymptote except in the case that all trials are eventually rewarded (*αC*_0_ = 1; Figure 2c; see Supplemental Information for mathematical details). Otherwise, greater willingness to wait increases expected time spent at the port on each trial (in the extreme case where no trials are rewarded, the expected time at the trial is equal to willingness to wait). Figure 2d shows how expected reward per trial increases as the expected time at the port increases when *t*_*w*_ is varied, combining the information in Figures 2b and c.

We can maximize the total reward rate in equation 5 by computing its derivative and setting it to zero, which yields

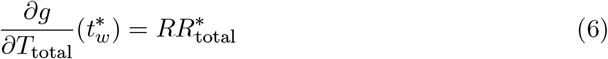

where 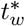 and *RR*^*∗*^ are the reward-rate-maximizing willingness to wait and the corresponding reward rate at that optimal 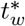 (see supplement for derivation). This is a generalization of the marginal value theorem^12^ to the case of stochastic rewards^15^. That is, equation 6 states that the prescribed rule for maximizing reward rate is to be willing to wait for reward until the time, 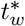, when the derivative of expected reward rate within the trial, 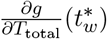, falls to the level of the maximum achievable reward rate across trials, 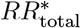. The latter quantity is the opportunity cost of continuing to wait for reward rather than beginning a new trial.

As noted above, we can simplify the computation by ignoring the reward consumption time in the denominator of equation 5. Instead, we will maximize the expected reward per time spent pursuing (not consuming) reward, *T* (*t*_*w*_):

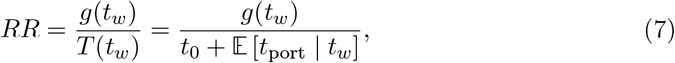

which is maximized when

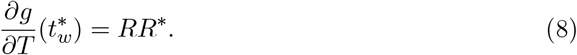

(Note that previous work^5^ assumed the optimality condition 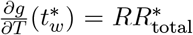, which produces suboptimal behavior when *t*_drink_ ≠ 0, as is the case in our data.)

This reward maximization rule can be understood graphically by plotting expected reward in a trial, *g*(*t*_*w*_), as a function of expected time pursuing reward in the trial, *T* (*t*_*w*_), for an example agent (Fig 2d,e). The reward rate for any choice of *t*_*w*_ will be equal to the slope of a line that passes from the origin through the point (*T* (*t*_*w*_), *g*(*t*_*w*_)). The maximum possible slope (i.e., maximum possible reward rate) is achieved when this line is tangent to the reward rate curve satisfying equation 8 (Fig 2e,f). In standard optimal foraging theory, the forager receives continuous reward and gives up and moves on at a time under its full control, 𝔼 [*t*_port_ | *t*_*w*_] = *t*_*w*_, which would mean that 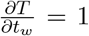 and equation 8 reduces to the marginal value theorem. However, because our task provides at most one reward per trial, the agent must estimate the expected rate of reward in each trial through experience and then set an upper bound, *t*_*w*_, on the time it will spend waiting for reward delivery.

### Optimizing wait time

Now that we have found the condition under which reward rate is maximized (equation 8), we are able to find 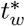 for a given set of environmental statistics. To do so, we first compute the derivative in left hand side of equation 8, which we write as

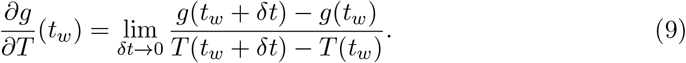

For an agent that has already waited for time *t*_*w*_, this quantity takes one of two values. If reward has already been delivered, the derivative is zero (*g*(*t*_*w*_) = *g*(*t*_*w*_ + *δt*) = 1), and the agent should move on to the next trial as soon as it finishes consuming reward. In the second case, the agent has not yet received the reward (*g*(*t*_*w*_) = 0). In this case, the expected reward after waiting for an additional time step is the probability of receiving reward in the next time step given that it hasn’t been delivered so far, *P* (*r*_*w*_ | ¬ *R*_*w*_). This quantity is the hazard rate of the distribution of reward delays including the trials in which no reward is baited. We refer to this quantity as the instantaneous reward expectation after waiting for time *t*_*w*_ without reward. We can write it as a product of the reward hazard rate for trials where reward is baited (equation 2) and the posterior probability that reward will be delivered in a trial given that it has not been delivered so far:

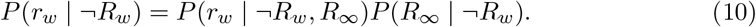

We refer to the second term as the agent’same posterior belief that reward will be delivered on a given trial after waiting for time *t*_*w*_ without receiving reward. We write this quantity using Bayes’ rule

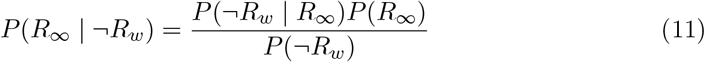

and evaluate it for the distribution used in our experiment

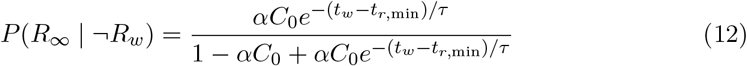

(see Supplemental Information materials for detailed derivation). This quantity has the value *αC*_0_ at the time of choice (*t*_*w*_ = 0) and falls to 0 as time passes. Note that when the agent is unaware of the probe trials (i.e., the agent estimates *α* = 1), equation 12 is equal to the posterior belief that the port choice was correct, the posterior decision confidence, after waiting for time *t*_*w*_ without reward.

Substituting equations 2 and 12 into equation 10, we get the instantaneous reward expectation after waiting for time *t*_*w*_ without receiving reward

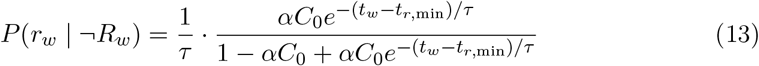

for *t*_*w*_ ≥ *t*_*r*,min_ (instantaneous reward expectation is 0 for *t*_*w*_ *< t*_*r*,min_). Note that equation 13 is equivalent to equation 5 in Lak et al.^5^ if we substitute *C* = *αC*_0_ and *t* = *t*_*w*_ − *t*_*r*,min_. However, our derivation clarifies that even though the reward hazard rate is fixed in the task, there is a decrease in instantaneous reward expectation while waiting for reward that can be attributed to a decrease in the posterior belief that the trial will be rewarded as time passes without reward delivery. Later, we will make use of this observation to develop a model for describing the port-leaving decision as a process that unfolds in time.

Instantaneous reward expectation (equation 13) for the example agent is plotted as a function of elapsed time without reward in Figure 2g. To find 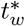, the agent needs to estimate the instantaneous reward expectation as a function of *t*_*w*_ and choose a moving on threshold, *κ*, whose optimal value is *RR*^*∗*^ (Fig 2g). When *κ < RR*^*∗*^, the agent is impatient and receives a below average reward rate, and when *κ > RR*^*∗*^, the agent wastes time at the reward port that would be better spent starting a new trial. Choosing the appropriate threshold will lead to optimal waiting with

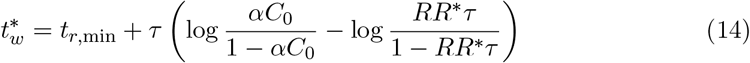

(see Supplementary Information for detailed derivation). This is equivalent to equation 6 in Lak et al.^5^ if we again substitute *C* = *αC*_0_, set *t*_*r*,min_ = 0, and substitute *κ* = *RR*^*∗*^. But note that in Lak et al.^5^, *κ* is defined in words as the “environmental reward rate” whereas we have developed an explicit expression for *RR*^*∗*^, which clarifies that the correct *RR*^*∗*^ in this expression is not the total environmental reward rate (equation 5), but rather the reward per time spent pursuing reward (equation 7). Moreover, because we have provided an expression for *RR* that makes all relevant environmental statistics explicit, we can now compute 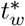 for a given experiment, which was not possible previously.

The optimal strategy for maximizing reward is influenced by the confidence on a given trial and all of the factors that influence the maximum achievable reward rate, including the probe trial fraction, the reward delivery time constant *τ*, and the travel time, *t*_0_. As *t*_0_ increases, the maximum possible reward rate decreases and the value of 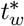 increases. The reward optimization procedure is depicted for three example levels of travel time in Figure 2h,i.

We found the optimal willingness to wait, 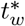, as a function of travel time, *t*_0_, for all the levels of *αC*_0_ by using root finding to solve equation 8 (Figure 2j; see Methods for details). The amount of time that a reward rate maximizing agent is willing to wait in this task increases monotonically as travel time increases for all levels of *αC*_0_ (except 0 and 1, where the agent should either not be willing to wait at all, or should always be willing to wait until the reward is delivered, respectively).

### Rats maximize reward rate after accounting for travel time

Now that we can compute 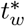 for an agent with fixed confidence across trials, we can test whether our rats achieved the maximum possible reward rate across trials. To do this, we compared rats’ willingness to wait, averaged across trials, to 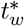, the willingness to wait that would maximize the reward rate for an agent with fixed confidence across trials. To estimate the rats’ average willingness to wait across trials, we computed the average wait time for correct probe trials and for a subset of error trials, subsampled so that the proportion of error trials used in this analysis matched the proportion in the full dataset (which also includes non-probe trials; Fig 3c,d). To compute 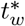 for each subject’s dataset, we estimated the necessary terms from that subject’s data: *α* was the fraction of non-probe trials in the rat’s dataset, *C*_0_ was the fraction of correct trials, and *t*_0_ was estimated from the mean travel time for the rat (after excluding the longest 1% of travel times, because the rats occasionally fully disengaged from the task for long periods of time; Fig 3a, b). Using these terms allowed us to evaluate both sides of equation 8 and compute 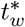 by root finding (see Methods for details).

**Figure 3:**
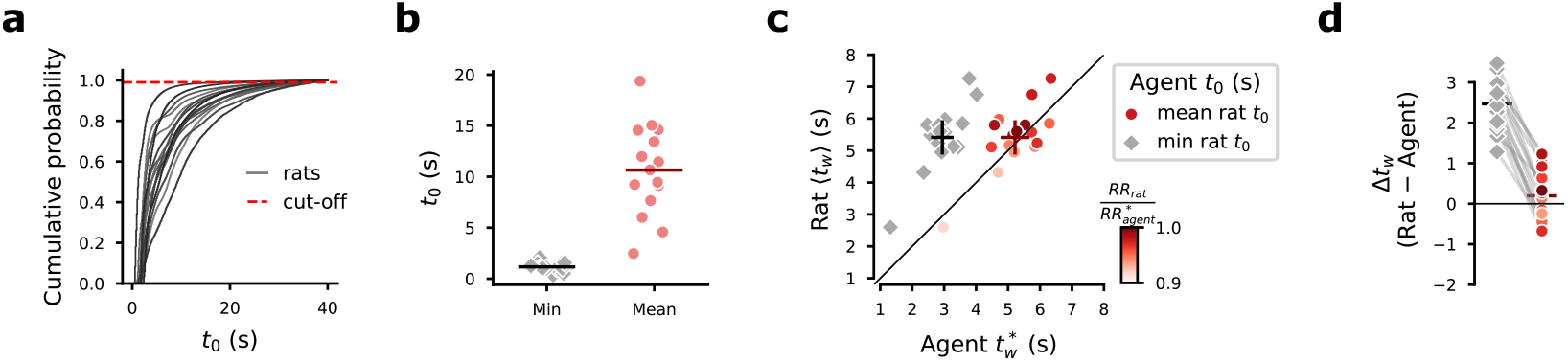
Rats maximize reward rate after accounting for travel times. (A) Cumulative distribution of travel times, *t*_0_, for each rat (black traces). We excluded the longest 1% of travel times for each rat to compute the mean travel time (dashed red line). (B) Minimum *t*_0_ for each rat (gray diamonds) and mean included *t*_0_ for each rat (red points). Means of the presented values are shown as horizontal lines for the minimum and mean *t*_0_ values. (C) Each rat’s willingness to wait is plotted as a function of the optimal willingness to wait for an agent with the same reward delivery statistics, trial accuracy, and the same mean travel time as the rat, but without trial-by-trial variations in confidence (red points). Each rat’s willingness to wait is also plotted as a function of the optimal willingness to wait for an agent as described, but with travel time equal to the minimum travel time achieved by the rat (gray diamonds). Rat willingness to wait is estimated by computing the mean wait time in correct probe trials and in a subsample of error trials (subsampled so that the proportion of error trials in this analysis is equal to the proportion of error trials for the full dataset when all correct trials are included). Optimal willingness to wait was determined by root finding using equation 8. The mean and 95% confidence intervals are shown as crosses for each group. The shade of the red points indicates the fraction of the reward maximizing agent’s reward rate achieved by the rat. For the comparison between the rat and the agent with matched mean *t*_0_, the shade of red of the points indicates the fraction of the maximized agent reward rate achieved by the rat. (D) Difference between the rat data and the agent data for the rats’ travel times (red points) and for the shorter travel times (gray diamonds). Colormap is the same as in C. The mean difference for each group is marked with a horizontal line.

A fully optimal agent performing this task should spend as little time traveling from one reward opportunity to the next. However, our rats spent more time than necessary traveling between reward opportunities. This was due to the self-paced nature of the task and exacerbated by center poke violations, which caused trials to be invalidated, further delaying time to the next reward opportunity. We reasoned that it may be very difficult for the rats to further minimize travel time. Among other things, decreasing travel times would require reducing the center port violation rate, which the animal is presumably already incentivized to do as much as possible. Therefore, we treated travel time as a constraint experienced by the animal and used the agents with matched travel times to ask whether the rats maximized reward rate given this constraint. To compare each rat to an agent who had also minimized travel time, we also computed 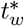 for an agent whose average *t*_0_ was set to the value of the shortest travel time achieved by the rat across sessions (Fig 3b).

We found that the rats’ average willingness to wait was not different from the reward maximizing agents’ with matched travel times (*p* = .25, paired t-test; Fig 3c,d). However, rats’ wait times were much longer than those of the agents optimized with short travel times (*p* = 6.15 × 10^*−*11^, paired t-test; Fig 3c,d). Subjects “overharvested” relative to fully optimal agents, as has been seen in previous studies of foraging behaviors^17,14^. However, when travel time is treated as constrained, and behavior is optimized over *t*_*w*_ alone, subjects’ overall willingness to wait was near-optimal, approximately maximizing their overall reward rate.

### Process model for optimal confidence-modulated waiting

To understand how the port-leaving decision might be implemented in the brain, we developed a process model that described the decision to stop waiting as an accumulation to bound process. This model provides us with a cognitively tractable algorithm that can achieve optimal waiting and model the cognitive state of the animal during this task, which may be useful for studies of neural recordings in the task. It also allows us to capture variability in waiting that may be explained by sources of internal variability other than variations in decision confidence.

To develop such a model, we used the sequential probability ratio test^18^ to derive a tractable update rule, a linear drift with time, for a decision variable that can be used to produce optimal wait times. From equations 8 and 10, we know that optimal policy is to stop waiting when the instantaneous reward expectation falls to the level of the maximum reward rate in the environment, or equivalently, when the posterior belief that reward will be delivered eventually, *P* (*R*_∞_ | ¬*R*_*w*_) falls to the maximum reward rate in the environment scaled by *τ*

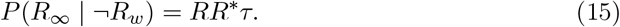

We can describe an agent who uses this strategy, but does not necessarily choose the optimal bound by replacing *RR*^*∗*^ with a parameter *κ* whose optimal value is *κ*^*∗*^ = *RR*^*∗*^. This process is shown in Figure 4a.

**Figure 4:**
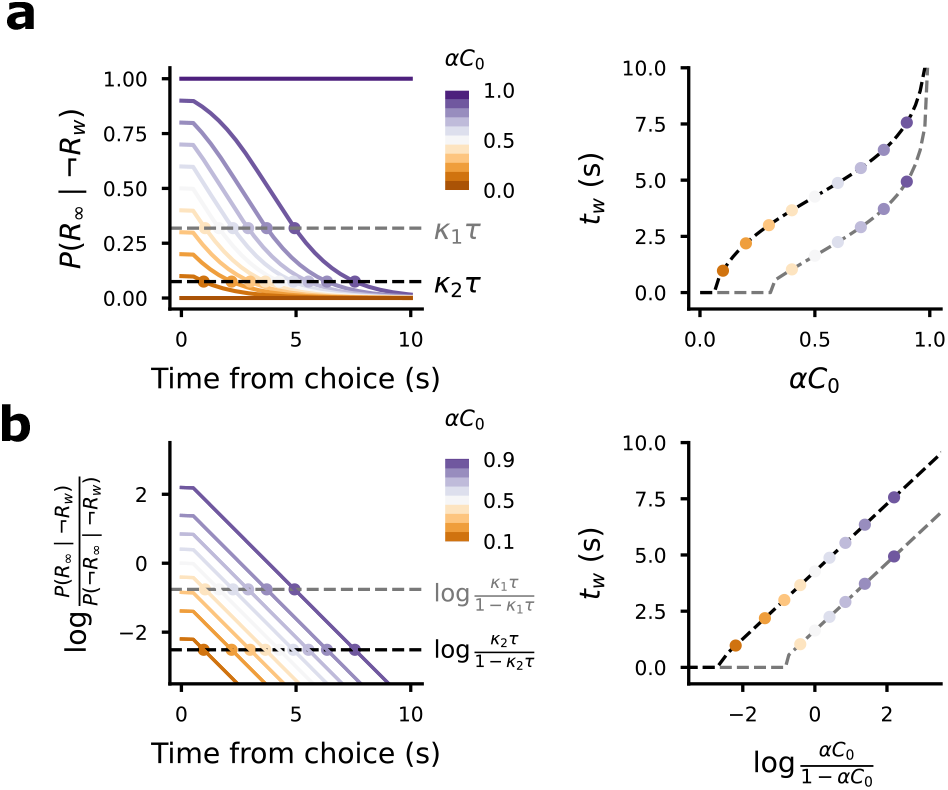
Optimal waiting can be implemented with linear drift from a confidence-dependent initial point to a fixed bound. (A) Optimal wait time procedure as presented in fig 2g,i, but scaled by *τ*, so that we track the evolution of *P* (*R*_∞_ |¬*R*_*w*_) to a bound *κτ*. Two settings of *κ* are shown (gray and black dashed traces) along with the bound hitting times for different levels of *αC*_0_ (left) and the resulting wait times are plotted against *αC*_0_ (right). (B) Equivalent model with linear drift from an initial point 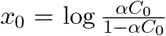, which terminates at a bound 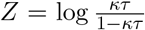. This process leads to the same waiting times as in A. Colormaps are the same as in A, but note that *C*_0_ = 0 and *C*_0_ = 1 do not appear in panel B, because they start at −∞ and + ∞, respectively, and never hit the bound.

To produce a decision variable that is tractable to update, we define *x*_*w*_ as the log odds of eventual reward delivery, given that reward has not been delivered by time *t*_*w*_:

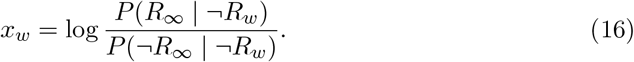

To find an update rule that integrates the information from the passage of time without reward into *x*_*w*_, we decompose *x*_*w*_ into two terms representing the previous value *x*_*w−*1_ and an update Δ*x* when timestep *w* elapses without reward (note that if reward is delivered in timestep *w*, the process ends):

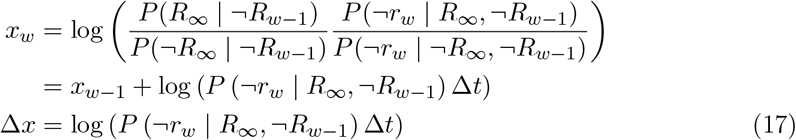

In the time before the earliest possible reward delivery (*t*_*w*_ *< t*_*r*,min_), the update term is 0 and *x*_*w*_ is constant, afterward *x* drifts at the hazard rate of the reward distribution

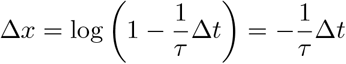

where we have used log(1 −*n*) ≈ −*n* for |*n*| *<<* 1. Taking the timestep to zero, we get the linear drift dynamics

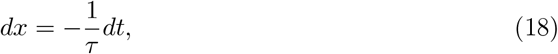

which we can combine with equation 4 to write *x*_*w*_ as a function of it’s initial value *x*_0_:

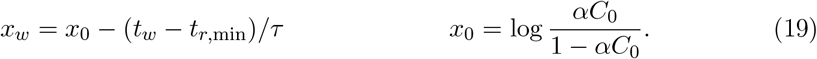

By equations 15 and 16, we know that the agent should stop waiting when *x*_*w*_ hits a bound specified by

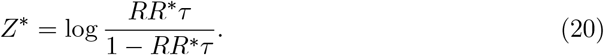

If the waiting process terminates when *x* hits the bound *Z*^*∗*^, we achieve the reward maximizing wait times equivalent to equation 14:

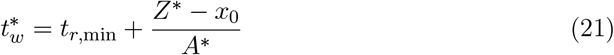

where *A* is a drift rate whose optimal value is *A*^*∗*^ = −1*/τ*. The evolution of *x*_*w*_ and equivalence of this waiting process with that of Figure 4a is shown in Figure 4b. This expression for optimal willingness to wait is equal to equation 6 in Lak et al.^5^ after setting *t*_*r*,min_ = 0 and *C* = *αC*_0_. But, now also has an algorithmic interpretation that may be possible to implement in the brain. In words, optimal wait time decisions can be made by initializing a decision variable at a value that is set by the decision confidence and evolving it toward a fixed bound that is set by the overall reward rate and reward delivery timing in the task. The drift rate toward that bound is set by the reward hazard rate.

### Contributions of different sources of timing noise to waiting process

In studies of timing judgments, subjects often exhibit the phenomenon of scale invariance in which the standard deviation of timing estimates increases linearly in proportion to the interval to be timed^21,22^. A previous model^5^ of confidence-guided waiting behavior assumed that scale invariant timing noise was the dominant source of noise affecting wait times. However, this has not been directly tested and it is not clear that timing in this task is dominated by the same sources of variability as in interval timing tasks where the goal is to learn to respond when reward is most likely, rather than to persist until the moment that reward is sufficiently unlikely that it is worth giving up and moving on. It is also not trivial to separate noise in the timing decision from variability in the confidence level that might result from any given percept.

We used the process model defined in the previous section to examine the patterns of timing variability expected when different aspects of the process were corrupted by noise. We considered three possible ways of adding timing noise to our process model. In the first, we add noise to the drift rate, *A* (Fig 5a). This produces scale invariant variability. For a given initial point, *x*_0_, the standard deviation in bound hitting times that is proportional to the mean. In the second model, we added diffusion noise to the position of *x* at each time step, which adds a random walk to the deterministic drift (Fig 5b). In this model, the standard deviation of wait times is proportional to the square root of the mean hitting times for a given *x*_0_, meaning that the standard deviation will grow slower than for the scale invariant model. Finally, we considered a model with a noisy bound (Fig 5c). In this model, the standard deviation is constant regardless of the initial point *x*_0_. The noise parameters were chosen for each model to produce a coefficient of variation (CV; ratio of standard deviation to the mean) of 0.3 when *x*_0_ = 0. For the scale invariant model, the CV is 0.3 for all values of *x*_0_, which is the level of noise assumed in Lak et al.^5^ .

**Figure 5:**
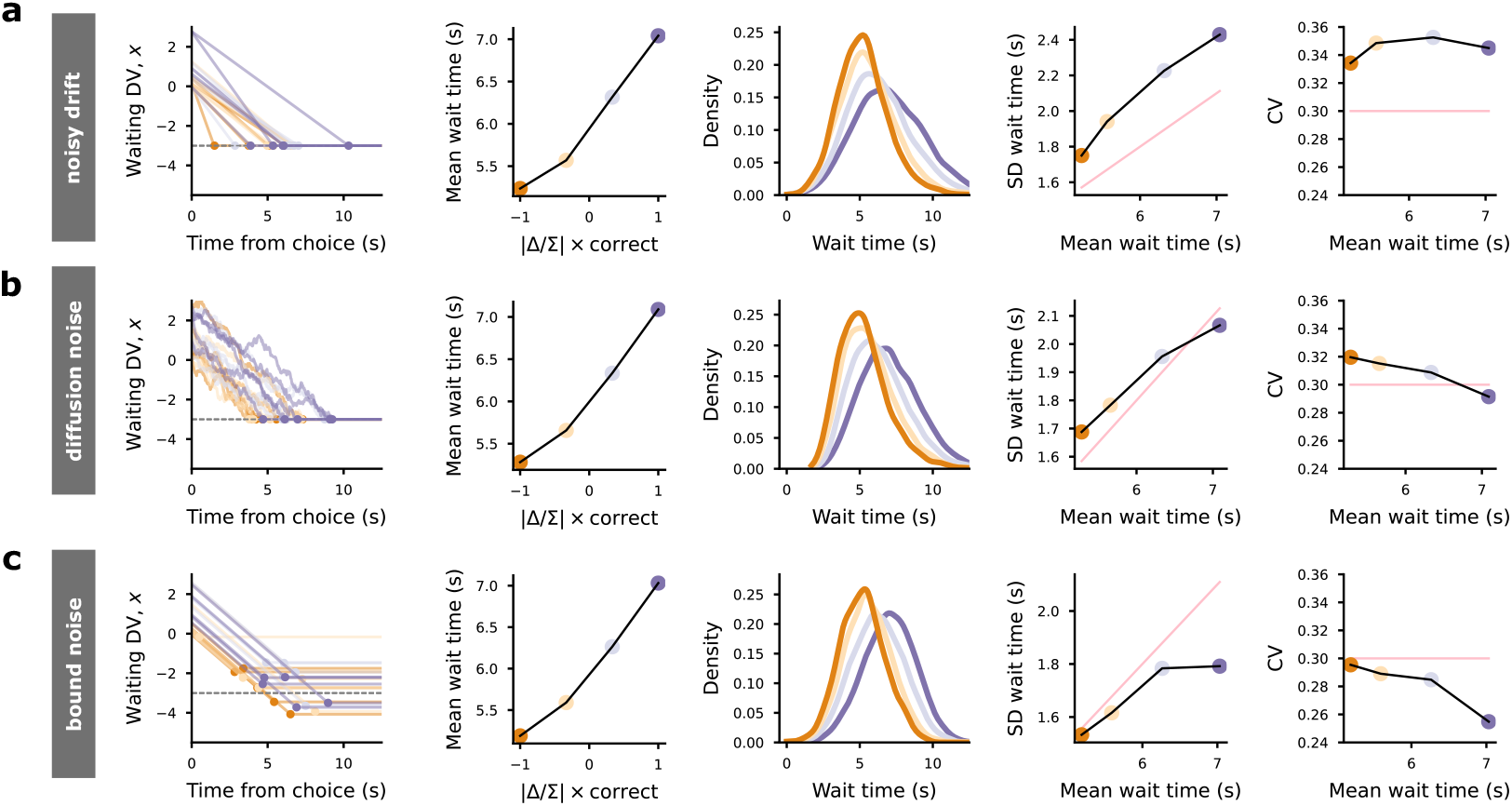
Candidate timing noise models. (A-C) We consider three candidate models for adding noise to the wait time decision. In all three models, the agent makes left/right port choice based on a stimulus, which is corrupted by perceptual noise and creates an accompanying decision confidence, *C*_0_. The waiting decision variable (DV), *x* is created by setting 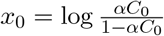 where *α* is the fraction of non-probe trials. And *x* drifts with rate *A* toward a boundary *Z*. The agent is willing to wait for reward until *x* hits the bound *Z*, but gives up and moves immediately when the bound is hit. (A) Left panel shows example particles from a model in which the noise comes from variability in the drift rate. This noisy drift model produces the scale invariant property in which the ratio of the standard deviation of hitting times to the mean of hitting times is constant across all *x*_0_ values. Traces are colored according to the binned normalized evidence favoring the choice, 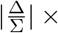 correct. Second panel from left shows kernel density estimates of bound hitting times for each of the bins of normalized evidence favoring the choice with the same colormap. The third panel from the left shows the mean wait time as a function of the normalized evidence favoring the choice (black trace with points colored by bin). The second panel from the right shows the standard deviation of the bound hitting times as a function of the mean in each bin (black trace with points colored by bin). The generative relationship between standard deviation and mean is underlaid (pink trace). Any deviation reflects noise added by the psychometric decision process. The rightmost panel shows the coefficient of variation (CV) as a function of the mean wait time in each bin (black trace with points colored by bin). Again, the generative relationship is underlaid (pink trace). (B) Plots are as in A, but for a model with a diffusion noise process in which noise is added in every time step. The pink traces from A are maintained for comparison. (C) Plots as in A and B, but for a model in which noise comes from variability in the bound. The pink traces from A and B are maintained for comparison.

While the patterns of timing variability produced by each of these models are simple when the initial wait time decision variable, *x*_0_, is known, we don’t have access to *x*_0_ for our rats. To understand the pattern of variability expected under each model when *x*_0_ is unknown, we generated simulated *x*_0_ values for 50,000 trials. To do this, we supposed a signal detection theory model of the decision process in which the stimulus is characterized by the ratio between the click difference and the total number of clicks on each trial, 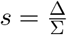. For each trial, we generated a percept by adding noise with standard deviation *σ*_*s*_ to the stimulus, *p* = *s* + *ξ* where 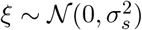. Decisions we made by comparing the stimulus to a decision boundary, *b* = 0. Confidence was then defined (beginning with equation 3) as

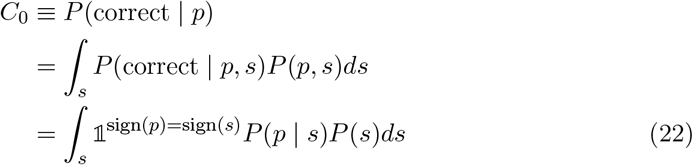

where we are integrating the probability of experiencing the percept *p* given the stimulus *s* over all levels of *s* that would produce a correct choice (see Supplemental Information for full equations). For the simulations, we assumed a uniform prior, *P* (*s*), and the probability of a percept given a stimulus is the Gaussian 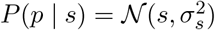. We used a value of 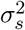 that best fit an example rat (see Methods for details). We assumed an accurate estimate of the non-probe trial frequency, *α*. Combining this with confidence, we produced a sample *x*_0_ for each trial using equation 19. We then generated a sample willingness to wait on each trial by applying the drift dynamics (equation 18) until the particle hit a bound *Z* (set to −3 in the simulations), chosen to produce a similar range of wait times as observed in data.

To determine what patterns we would be able to see in our rat data, we analyzed the simulated dataset for models with each source of timing noise as though we did not know generative *x*_0_, but could only observe the stimulus, choice accuracy, and willingness to wait on a given trial. We binned trials by the evidence supporting choice, 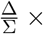 correct. We then plotted conditioned kernel density estimates of willingness to wait for each bin. We also computed the mean, standard deviation, and coefficient of variation (standard deviation divided by mean) in each bin.

All models achieved increasing mean willingness to wait as a function of evidence supporting choice. But, the models had different relationships between standard deviation and mean within each bin. The variable drift model produced a roughly proportional increase in standard deviation as the mean grew, corresponding to flat coefficient of variation, as expected under scale invariant noise (Fig 5a). But, there was additional noise across all bins that arose from variability in confidence (*x*_0_) within each bin. The diffusion noise model produced a slightly sublinear increase in standard deviation as the mean grew, corresponding to a decreasing coefficient of variation (Fig 5b). Finally, the bound noise model produced a distinctly sublinear increase in standard deviation as the mean grew, with almost no increase between the last two evidence bins (Fig 5c). This corresponded to a dramatic decline in the coefficient of variation for the simulated data.

### Scalar variability is not the dominant source of noise in rat waiting data

To test whether scale invariant timing noise was the dominant noise source affecting rats’ waiting time decisions, we analyzed the rat data using the analysis methods used to study the simulated data. We analyzed the mean and standard deviation of probe trial wait times in bins of normalized evidence strength favoring the rat’s choice (Δ*/*Σ × correct). The average wait time in each bin is shown for an example rat in Figure 6a. The distribution of wait times in each of these bins is shown for the example rat in Figure 6b. We compared the standard deviation of wait times in each of the bins to the mean (Fig 6c) and computed the coefficient of variation in each bin (Fig 6d) for the example rat. The pattern we observed was not consistent with the simulated data for the model with scale invariant timing variability caused by noisy drift. Instead, the pattern we observed was sublinear, with minimal increase in standard deviation between the last two bins.

**Figure 6:**
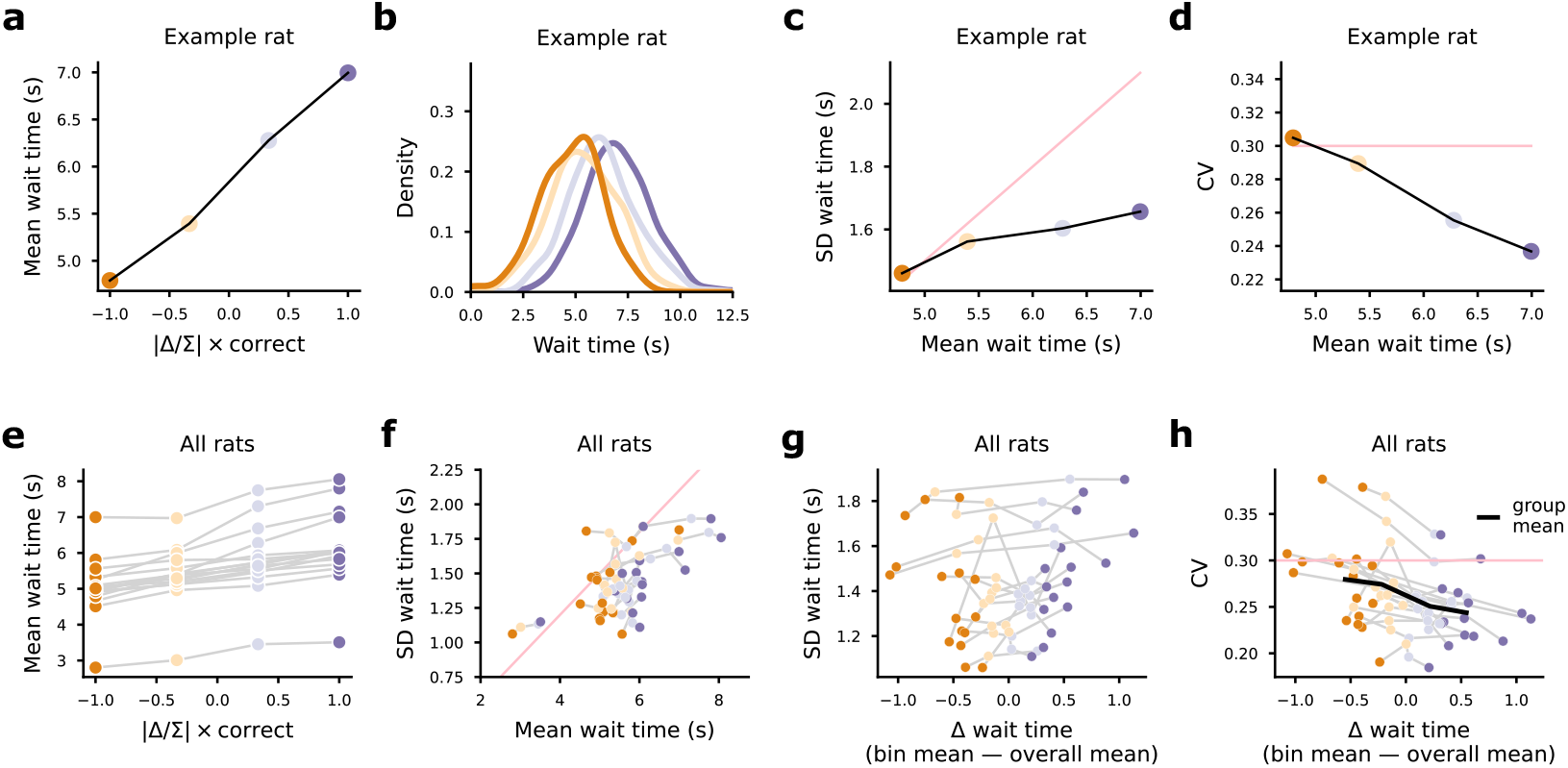
Scale invariant noise is not the dominant source of variability in the data. (A) Average wait times for an example rat for correct probe trials and proportionally sampled error trials as a function of evidence for chosen option bin. (B) Kernel density estimate for the wait times in each bin from A (the same colormap is used to indicate evidence bin). (C) Standard deviation of the wait times in each bin (points connected by black trace) is overlaid on the predicted relationship between wait time standard deviation and mean under scale invariance with a coefficient of variation equal to .3 (pink trace). (D) Coefficient of variation (the standard deviation divided by the mean) in each bin is plotted as a function of mean wait time in each bin for the data (points) and for the scale invariant model shown in C (pink trace). (E) All rats’ mean wait times plotted as a function of evidence for chosen option as in A. (F) All rats’ wait time standard deviations plotted against mean wait time as in C. Again, overlaid on the prediction from scale invariance. (G) Standard deviation in each bin for all rats plotted against the difference between the mean wait time in the bin and the average of the bins for that rat. (H) Coefficient of variation plotted as in G for all rats with the mean across rats overlaid (black trace). If scale invariance was the dominant source of noise, each rat’s trace should be flat.

We repeated this procedure for all rats. Average wait time as a function of binned evidence for choice is shown for all rats in Figure 6e. The standard deviations as a function of average wait time are plotted for all rats in Figure 6f (as in Figure 6c). To compare across rats, we subtracted off the average wait time of the bins from each rat’s data (Fig 6g). We find a consistent pattern across rats that the relationship between standard deviation and mean wait time is flatter than expected under the scale invariant model. We then examined the coefficients of variation along this axis and plotted them together for comparison (Fig 6h). Across rats, we see a consistent downward trend in the coefficients of variation, inconsistent with scale invariant timing noise. This suggests that other sources of noise dominate any scale invariant noise that exists in our rats’ behavior. Additionally, the standard deviation appears to increase more slowly than expected if diffusion process was the dominant source of noise. Qualitatively, the variability in our data appears most consistent with the model in which variability in the bound dominates. This variability may stem from noise, but may also stem from continual learning of the appropriate bound setting as a recency-weighted average of the reward rate history^11,14^.

## Discussion

We trained rats to perform a task requiring auditory evidence accumulation^16^ combined with a post-decision temporal wager designed to assess their decision confidence^5^. The time that animals are willing to wait for a delayed reward after making a decision has become a popular proxy measurement of decision confidence, because we know that optimal agents wait longer for rewards they are more confident they will receive^5,7,10^. However, willingness to wait in optimal agents is also influenced by the maximum possible reward rate in the environment, which is in turn influenced by many environmental statistics. These statistics determine the optimal overall average willingness to wait, but they are not explicitly accounted for in previous studies of confidence-guided waiting.

Here, we developed an expression for the reward rate in the environment which made all of the relevant environmental statistics explicit. Using this environmental reward rate, we derived an expression for the conditions under which reward was maximized in the environment. This generalized the marginal value theorem^12^ into the case of stochastic rewards^15^ with arbitrary initial expectations about the probability of eventual reward. This work made it possible to test whether rats performing this task achieved overall reward-rate-maximization, which we refer to as “optimal” behavior.

One of the key statistics that determines the optimal overall average willingness to wait is the travel time incurred when deciding to move on from a given reward opportunity and pursue the next. We observed that our animals were willing to wait longer than fully optimal agents who minimized travel time and then maximized reward rate by finding the best overall willingness to wait for that minimized travel time. Compared to these agents, our animals “overharvested” reward on each trial, as has been seen in other studies of foraging behaviors^17,14^. Unlike fully optimal agents, our animals took longer to travel between reward opportunities than was strictly necessary. We asked whether their waiting behavior was optimal if we treated their travel times as constrained, meaning that they had minimized travel time to the best of their ability and then optimized willingness to wait given those travel times. We found that when we treated travel times as constrained, the rats’ waiting behavior was near optimal.

One limitation of our study is that we don’t know for certain that our rats’ have minimized travel time to the best of their ability. It is possible that rats could decrease time between trials and achieve a higher reward rate. One way that future studies could test this would be to impose a longer minimum intertrial interval and test whether rats found a new willingness to wait that was optimal for the increased travel times.

To understand how our rats achieved near-optimal wait times, we developed a model of the wait time decision process, which sought to capture the unfolding cognitive state throughout the decision. Taking inspiration from the success of the drift diffusion model in modeling two alternative decisions^27,20,24^, we used the sequential probability ratio test^18^ to develop an optimal update rule for a decision variable that can control the port-leaving decision. This produced a continuously evolving cognitive process model for controlling port-leaving time via the linear drift of a decision variable toward a fixed bound. This process model provided a tractable algorithm for implementing optimal waiting in the brain in which each of the separate parameters could be learned from experience.

The model also allowed us to consider sources of variability that might contribute to the decision process, drawing on extensive work on pacemaker accumulator models of timing behavior^22^. To understand the mapping between willingness to wait and confidence, it is useful to know what sources of variability are contributing to wait times. Previous work has assumed that variability in the timing of willingness to wait would be dominated by the scale invariant property^5^ in which the standard deviation of observed wait times should be proportional to the animals’ desired wait time^21^. However, this assumption had not been tested. We compared three models of timing variability in the waiting decision process. The first was a noisy drift model, which produced scale invariant timing noise. The second was a diffusion noise model, in which timing noise grew like the square root of the interval to be timed. And finally, a noisy bound model, in which timing noise was constant across desired wait times. We found that our data was most consistent with the model dominated by bound variability.

While our model provides an improved description of the port-leaving decision process, there are several avenues of possible improvement to the model that we should consider in the future. First, there are well-documented aspects of the port choice decision process that are not being accounted for here, including it’s evolution in time^16^, effects of trial history^28^, and change in the parameters of the decision process from trial to trial^29,3^. Second, there may be postprocessing of the stimulus following choice that leads to evolution of confidence independently from that instructed by the environment^30^. Finally, just as the parameters governing the choice process may evolve from trial to trial, the same may happen for the wait time decision either due to learning or changes in internal state like increasing satiety or patience^31^. Indeed, we speculate that continual learning of the bound controlling port-leaving may explain the variability we observed in our data.

Our model provides a tractable algorithm for solving this task, which can produce optimal behavior. The model can also produce a variety of forms of variability and deviation from optimality, which we have used to better understand the sources of variability in confidence-guided waiting decisions. Future work investigating the neural basis of confidence computations using the confidence-guided waiting paradigm should seek to link neural activity and perturbations of brain regions to the parameters of a dynamic model of the internal cognitive process for deciding when to give up and move on, like the one developed here. Using such a model will increase the interpretive power of experiments using this paradigm to understand how the brain computes confidence estimates and uses them to guide subsequent behavior.

## Methods

### Subjects

Animal use procedures were approved by the Princeton University Institutional Animal Care and Use Committee and carried out in accordance with NIH standards. All subjects were adult male Long Evans rats bred either at Princeton Neuroscience Institute (VGAT-ReaChR rats) or by one of the following vendors (wild type rats): Taconic, Hilltop and Harlan, USA. Rats were pair-housed unless implanted with infusion cannulae at which point they were single-housed. Rats were placed on a water restriction schedule to motivate them to perform the task for water rewards.

### Behavioral tasks

#### Poisson Clicks

We trained rats on the Poisson Clicks task^16^ with a post-decision wait time wager^5,6^ using an automated training protocol. Throughout training, rats were put on a controlled water schedule where they received at least 3% of their weight every day. Rats trained each day in training sessions of around 120 minutes.

In the final stage of training, each trial began with the illumination of a center nose port by an LED light inside the port. This LED indicated that the rat could initiate a trial by placing its nose into the center port. Rats were required to keep their nose in the center port (“center fixation”) for a fixed duration until the LED turned off as a “go” signal. During center fixation, two trains of randomly-timed auditory clicks were played from speakers on either side of the center port after a variable delay. The duration of the click trains was uniformly distributed. The two click trains were each associated with one of two side ports and clicks in each click train were generated using different Poisson rates. For a given rat, the two generative rates always summed to a fixed value (20 or 40 clicks *s*^*−*1^).

After the “go” signal, rats made a port choice by poking their nose into one of the two side ports. If they exited from the center port before the “go” signal, the trial was considered a violation and they experienced a white noise stimulus followed by a short time out. These trials did not yield decisions or wait times, but did contribute to travel times.

Choices were considered correct, and potentially rewarded, if they corresponded to the click train with the greater number of clicks, which corresponds to a noiseless ideal observer’s estimate of the larger click rate.

#### Confidence-guided waiting

Rewards were only delivered if the rat stayed at the side port until a reward time *t*_*r*_ drawn from an exponential distribution between a minimum *t*_*r*,min_ ∈ (.05*s*, .5*s*) and maximum *t*_*r*,*max*_ *>* 15*s* with time constant *τ* = 1.5*s*. The resulting mean reward delay was ⟨*t*_*r*_⟩ = *t*_*r*,min_ + *τ*. After errors, no feedback was delivered. Instead, the animal had to eventually give up on waiting for reward and start a new trial.

With probability *ζ* ∈ (.05, .15), the trial was turned into a probe trial by setting *t*_*r*_ = 100s. We did not allow multiple probe trials to occur consecutively. These probe trials allowed us to observe port-leaving times on a subset of correct trials when they might otherwise have been censored by reward delivery. Rats were given a grace period between 500 and 1500ms for leaving and returning to the choice port. If they withdrew from the reward port for longer than this grace period, reward was no longer available. If the rat returned to the center port, during or after the grace period, a new trial was immediately initiated. If they returned to the chosen side port after the grace period, or entered the opposite side port at any time, the possibility of reward delivery was removed. For analysis of uncensored wait times, we focused on trials where the rat initiated a new trial by center poking within 2 seconds of leaving the side port.

#### Shaping

We shaped the animals by first training them to perform the Poisson Clicks task via a standardized set of training stages. We then added the reward delay component. First, fixed feedback delays were introduced on both correct and error trials and grew in each trial until they reached *t*_*r*,min_. Then, the error feedback delay was incremented from trial to trial until the rat never waited long enough to get the error feedback. At that point, the error feedback delay was set to 100s. Next, the reward delays were randomized by gradually increasing the exponential time constant *τ* and the maximum delay time *t*_*r*,*max*_. When the *t*_*r*,*max*_ was larger than the rat’s longest waiting times, we set it to 100s. When *τ* reached it’s target value, we introduced probe trials. We did not allow multiple probe trials to occur in a row.

#### Inclusion criteria

Rats trained on this task were included in this study if they had more than 30 sessions that met the session inclusion criteria and if the fraction of unrewarded trials that ended with a re-initiating center poke (as opposed to re-entry in the chosen side port or entry into the opposite side port) met a minimum threshold of 55%. The session inclusion criteria required that the rat perform at least 150 trials with an overall accuracy rate exceeding 60%. In order to prevent the rats from developing biases towards particular side ports, an anti-biasing algorithm detected biases and probabilistically generated trials with the correct answer on the non-favored side.

#### Psychometric curves

Behavioral sensitivity was assessed using psychometric curves. The probability of choosing the rightward port was computed as a function of the binned normalized click difference 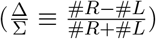. We fit psychometric curves with 2 parameters, a bias parameter *b* and a noise parameter *σ*, for all rats as a function of the normalized click difference. We fit the data by minimizing the negative log likelihood across trials where the probability of a rightward choice on a given trial was given by

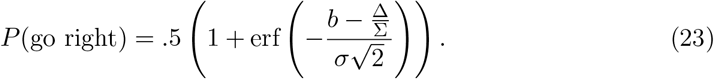

### Wait time chronometric curves

Wait time modulation was assessed using error trials and correct probe trials to create wait time chronometric curves, which relate mean wait time to the strength of evidence supporting the chosen option. Strength of evidence supporting choice was computed as 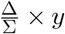, where *y* = ±1 with positive values for correct port choices and negative values for incorrect port choices. The trials with the most evidence supporting the chosen option have large, positive values and the trials with the most evidence against the chosen option have large, negative values. The most difficult trials, with the least evidence weighing on the choice, have small magnitudes. We expect confidence to increase monotonically along this axis. We fit a line to each rat’s wait times in the space of normalized click difference supporting the choice.

### Optimality analysis

To test whether rats’ waiting times maximized overall reward rate, we found the optimal overall average wait time, 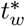, by evaluating equation 14 for each rat. To do this, we estimated the relevant terms contributing to equation 14 from the rats’ datasets: *α* was the fraction of non-probe trials in the rat’s dataset, *C*_0_ was the fraction of correct trials, *τ* and *t*_*r*,min_ were estimated from the reward delays scheduled for the rat, and *t*_0_ was estimated from either the mean travel time achieved by the rat (after excluding the longest 1% of travel times, because the rats occasionally fully disengaged from the task for long periods of time), or the minimum travel time achieved by the rat. Because we are only interested in average overall waiting time here, we don’t need to consider the variations in wait time associated with confidence. Therefore, this agent was constrained to wait the same amount of time on every trial, which allowed us to avoid making choices about how to capture variations in confidence for this analysis. We used a root finding algorithm to evaluate 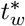 for a given set of task statistics. We compared the willingness to wait for each of these agents to the average waiting times for the corresponding rat in correct probe trials and in a subset of error trials (subsampled to ensure that the frequency of error trials in this comparison matched that in the overall dataset). We also measured the optimal agent’s reward rate and the fraction of the agent’s reward rate achieved by the rat.

### Process model simulations with candidate noise sources

We used euler integration to simulate the wait time decision process for three candidate noise models. In all simulations, we sampled 50,000 trial stimulus strengths, *s*, with replacement from the dataset of an example rat. We then generated a percept, *p* = *s* + *ξ*, for each trial, by adding Gaussian noise, 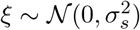, to the stimulus. The model made a rightward choice if the resulting percept was greater than a decision boundary, which we set to zero (i.e., *p > b* for *b* = 0). Given this percept, we generated confidence levels according to equation 22. We then produced a corresponding *x*_0_ and updated it in 25 millisecond timesteps (Δ*t* = .025*s*) according to equation 19. The drift was set to it’s optimal setting 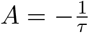 (per equation 18) and the bound was set to *Z* = − 3 to produce mean wait times across trials that roughly matched the example rat’s. To produce a model with scale invariant timing noise, and specifically a coefficient of variation of 0.3, as in Lak et al.^5^, we set the drift on each trial to be 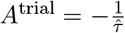 where 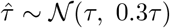. To produce a model with diffusion noise, we added Gaussian noise in each time step drawn from 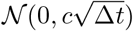 with 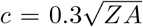 to produce an equivalent level of noise at *x*_0_ = 0 as produced under the scale invariant model. To produce a model with constant noise, we sampled a different bound on each trial *Z*^trial^ ∼ 𝒩 (*Z*, 0.3· |*Z*|). The magnitude of noise was again chosen to produce the same noise level as the other models for *x*_0_ = 0. We recorded each models willingness to wait on each trial as the timestep in which the particle *x* first crossed the bound *Z*.

### Analysis of variability in simulations and rat data

We used our simulations to ask what patterns of variability would be expected as a function of the stimulus. This was useful for analyzing rat data in which the confidence level and *x*_0_ level are unknown. To do this, we binned trials by the evidence supporting the chosen option for both the simulated data and the rat data. Within these bins, we computed kernel density estimates of the distribution, as well as computing the mean, standard deviation, and coefficient of variation (ratio of standard deviation to mean) of the wait times in each bin. These produced distinct patterns for each of the candidate models, which we then compared qualitatively to the rat data. In particular, the assumption of scale invariance predicted a flat coefficient of variation, which we did not observe in the rat data. Instead, our data was most consistent with the constant bound noise in which the standard deviation in each bin grows slowly as mean wait time increases.

## Acknowledgements

We thank Athena Akrami, Adrian Bondy, Diksha Gupta, Thomas Luo, all other members of the Brody lab, as well as Lukas Braun, Nathaniel Daw, Javier Masís, Stefano Sarao Mannelli, and Pat Simen for helpful conversations and suggestions. TB acknowledges support by NIH grant T32 MH 65214-16. This work was supported by a grant from the Simons Foundation (Grant # 542953) awarded to CB, as well as NIH grant R01MH108358 awarded to CB.

## Author contributions

T.B. and C.K. developed the rat training protocol. T.B. managed rat training and care.

T.B. and A.P. derived the equations and models. T.B. analyzed the data. T.B., A.P. and C.B. wrote the manuscript. C.B. oversaw all aspects of the project.

## Supplementary Information

### 1 Derivation of reward-rate-maximizing behavior

The total reward rate in the task is defined as the expected reward per trial, *g*(*t*_*w*_), divided by the expected time spent in each trial, *T*_total_:

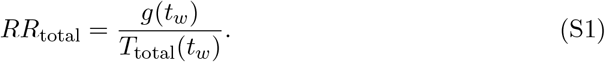

To maximize the reward rate, we find the condition such that its derivative is zero, 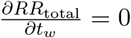. We compute the derivative using the quotient rule

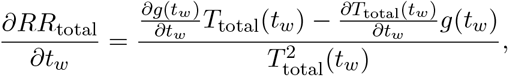

which is equal to zero when

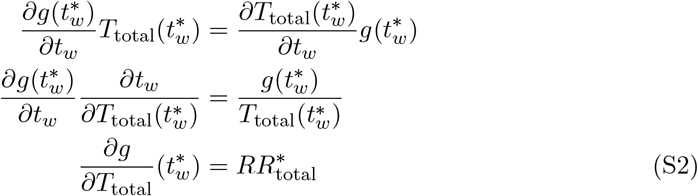

where 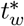 is the reward-maximizing willingness to wait and 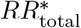 is the reward achieved at 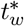 .

To find a solution for 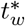 for a given set of task statistics from equation S2, we need expressions for *g*(*t*_*w*_) and *T*_total_(*t*_*w*_). To do so, we will use the notation introduced in the main text to simplify these expressions. We will use

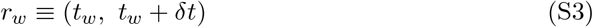

to indicate whether reward is set to be delivered in some infinitesimal timestep *δt* beginning at time *t*_*w*_. Then, we can indicate whether reward is set to be delivered before time *t*_*w*_ using the sum

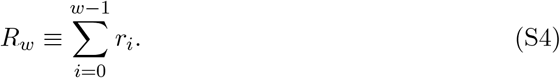

We will use the negation, ¬*R*_*w*_, to indicate when no reward is delivered by time *t*_*w*_.

#### 1.1 Expected reward per trial

Because at most 1 reward is delivered per trial and it always has the same magnitude, we can set the reward magnitude to 1 and make the expected reward per trial equivalent to the probability of reward in a trial

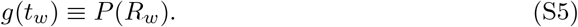

Our reward distribution is exponential, meaning that, given that reward is coming, the delivery times, *t*_*r*_, are distributed according to

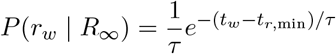

where *τ* is an experimenter-specified time constant and *t*_*r*,min_ is an experimenter-specified minimum reward time (equation 1 in the main text). The probability of receiving a reward before time *t*_*w*_ depends on this distribution and on the probability that reward will be delivered on this trial, *P* (*R*_∞_). The probability that reward will be delivered on this trial is estimated based on the decision confidence, *C* _0_, and the probability that a trial is not a probe a trial, *α*:

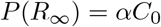

(equation 4 in the main text). The expected reward per trial as the probability of receiving a reward before time *t*_*w*_ is the cumulative density function for the exponential given that the reward is coming times the prior probability that the reward is coming, *P* (*R*_∞_):

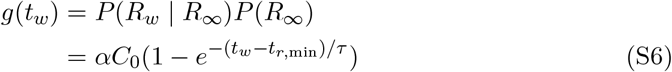

where we have used the cumulative density for an exponential to compute *P* (*R*_*w*_ | *R*_∞_).

#### 1.2 Expected time per trial

The expected time per trial can be broken into three epochs: the time between leaving the reward port on the previous trial and entering a reward port on the current trial, *t*_0_, the time spent at the port on the current trial, *t*_port_, and the time spent consuming reward on the current trial, *t*_drink_. Adding together the expected duration of each epoch, we get:

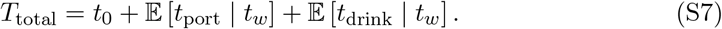

The first quantity is referred to as the “ travel time” and, for the reward-maximizing agent, does not depend on *t*_*w*_. The other two quantities depend on whether reward is set to be delivered and how long the agent is willing to wait. The consumption time, *t*_drink_, is either 0, if no reward is received, or a constant, if reward is delivered. Its expectation can be written

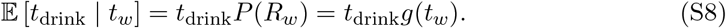

We will show that 𝔼 [*t*_drink_ | *t*_*w*_] can be ignored for the reward maximization process.

##### Expected time at the port

To compute the expected time at the port, 𝔼 [*t*_port_ | *t*_*w*_], we will separately consider trials in which reward is not set to be delivered and trials in which reward will be delivered if the agent waits long enough. Marginalizing over these possibilities gives us

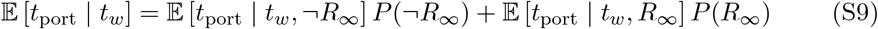

When no reward is set to be delivered, the agent always gives up and moves on at the time *t*_*w*_:

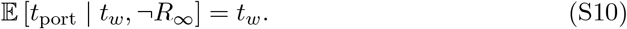

In trials where reward is set to be delivered at some time *t*_*r*_, the time at the port can take one of two values. If the agent is not willing to wait long enough to get the reward, the agent will give up before the reward is delivered and we will again observe time *t*_*w*_ spent at the port:

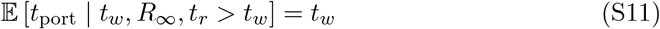

If the agent is willing to wait long enough to get the reward, the reward delivery will censor the agent’s willingness to wait and we will observe *t*_*r*_ time spent at the port. To compute the expected port time for this trial type, we need to marginalize over the possible values that *t*_*r*_ can take, as follows:

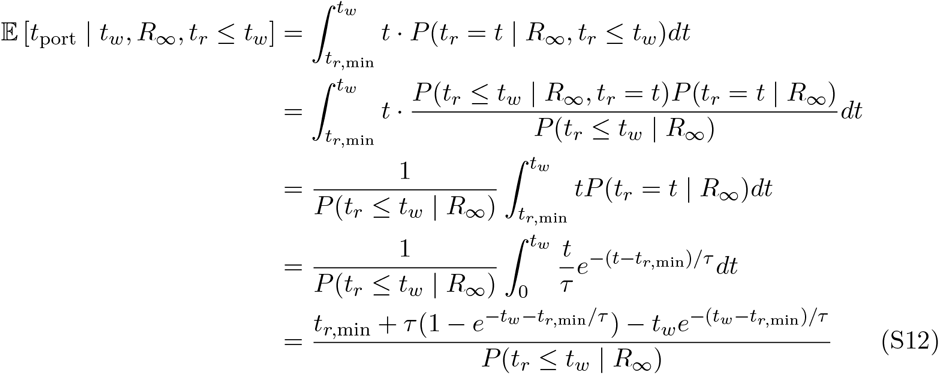

Combining equations S11 and S12 and multiplying each by their probabilities, we can compute the expected time at the port on trials where reward is set to be delivered eventually:

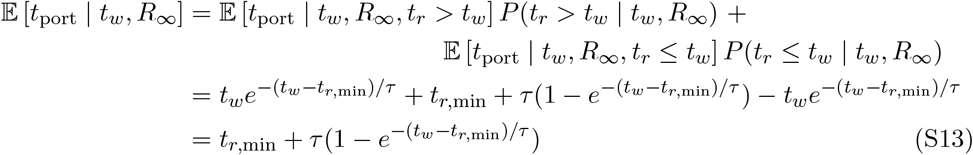

Finally, we can combine the expected port time in trials where no reward is baited (equation S10) and trials where reward is set to be delivered if the agent waits long enough (equation S13) to get the expected time at the port overall:

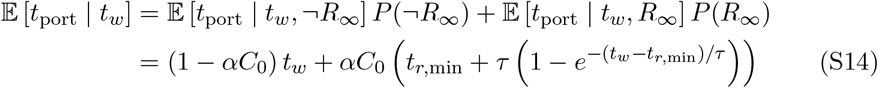

###### 1.2.1 Reward maximization doesn’t depend on consumption time

As mentioned above, we can ignore the consumption time in the reward maximization process, which simplifies equation S2. We will use *T* (*t*_*w*_) ≡ *t*_0_ +𝔼 [*t*_port_ | *t*_*w*_], to represent the expected time spent searching for, but not consuming reward. We will use 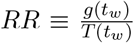 to represent the reward rate per time spent searching for reward. We can rewrite equation S2 with the consumption times made explicit and show that it can be ignored

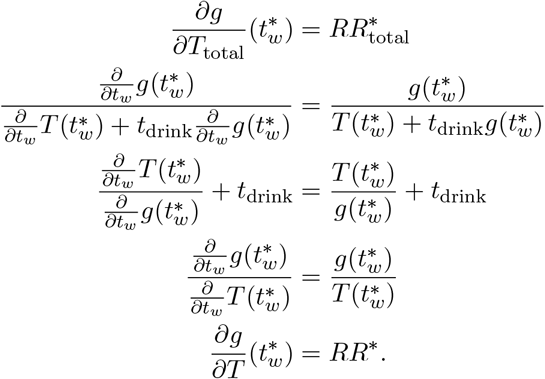

We will use equation 8 to find the optimal waiting behavior.

#### 1.3 Derivation of posterior belief that reward will be delivered

From equation 11 in the main text, we know that the posterior belief that reward will be delivered on a given trial after waiting for time *t*_*w*_ without receiving reward is

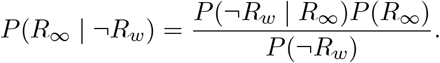

The first term in the numerator is the probability that reward is not delivered by time *t*_*w*_ given that it will be delivered eventually, which is the survivor function of the exponential distribution (or 1 minus the CDF)

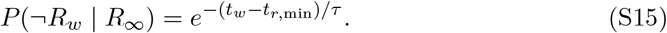

The denominator can be expressed as

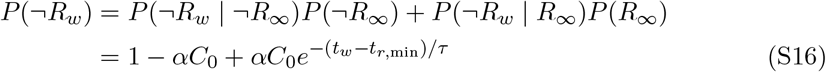

where we have used equation S15 and the fact that *P* (¬*R*_∞_) = 1 − *P* (*R*_∞_). Combining these expressions with the definition of *P* (*R*_∞_) (equation 4), we get:

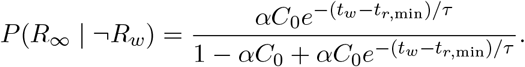

#### 1.4 Derivation of optimal willingness to wait

We rearrange the terms of the optimality condition from equation 8 and use the expression we derived for the instantaneous reward expectation (equation 13) to find the optimal willingness to wait

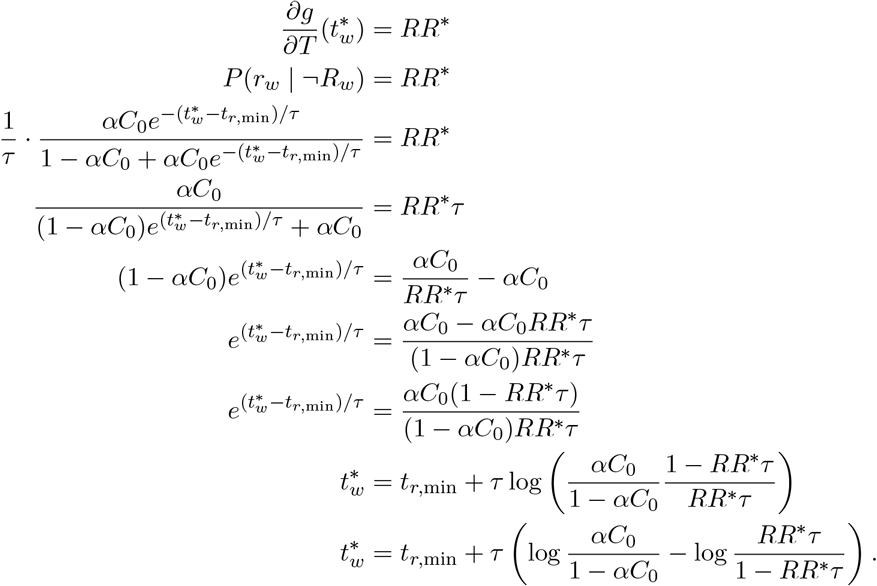

